# Optical imaging of treatment-naïve human NSCLC reveals changes associated with metastatic recurrence

**DOI:** 10.1101/2024.10.14.618213

**Authors:** Paola Monterroso Diaz, Jesse D. Ivers, Stephanie Byrum, Charles M. Quick, Konstantinos Arnaoutakis, Kyle P. Quinn, Narasimhan Rajaram

## Abstract

Lung cancer remains the leading cause of cancer deaths, comprising nearly 25% of all cancer deaths [1]. NSCLC accounts for approximately 85% of all cases and encompasses major subtypes such as adenocarcinoma and squamous cell carcinoma. Despite advances in surgical and therapeutic options, NSCLC remains associated with poor prognosis due to a high rate of recurrence, even in early stages. Around 30-55% of patients who undergo complete resection will experience metastatic recurrence, significantly lowering survival outcomes [2]. There is a critical need to develop prognostic markers capable of predicting risk of recurrence at earlier timepoints in order to improve NSCLC management, as it could help clinicians tailor treatment plans, optimize follow-up schedules, and identify high-risk patients who might benefit from adjuvant therapies. Two photon microscopy (TPM) techniques provide non-invasive high-resolution information on cell metabolism within tissue by utilizing an optical redox ratio (ORR) of FAD/[NADH+FAD] autofluorescence. The goal of this study is to use the ORR and NADH fluorescence lifetime decay to identify measurable differences in optical endpoints of human NSCLC that are indicative of their long-term outcome. Twenty-five treatment-naïve NSCLC specimens were classified into metastatic and non-metastatic groups according to subject-detail reports. The ORR and mean NADH lifetime were determined for each sample, revealing a significant increase in the ORR for the metastatic group. Additionally, tumors presenting with high optical redox ratios were found to be correlated with low KEAP1, a prognostic indicator of poor clinical outcome in NSCLC. To evaluate the prognostic potential of optical metabolic endpoints, we trained three classifiers: logistic regression, SVM, and KNN on three different feature sets: optical endpoints, clinicopathological features, and combination of optical and clinical features. We found that SVM trained on optical endpoints alone (AUC = 0.74) outperformed the model built with only clinical features (AUC = 0.62), when classifying tumors based on their metastatic recurrence status. Together, these findings highlight the potential of optical metabolic imaging to provide markers of recurrence in NSCLC.

## Introduction

Lung cancer remains the leading cause of cancer deaths worldwide, responsible for nearly 1.8 million cancer deaths each year [3]. Non-small cell lung cancer (NSCLC) accounts for the majority of diagnosed lung cancers (∼85%) and is most commonly classified into adenocarcinoma (ADENO) and squamous cell carcinoma (SCC) histological subtypes [4]. Due to the insidious nature and initially asymptomatic presentation of NSCLC, approximately 70% of NSCLC patients present with locally advanced or metastatic disease at time of diagnosis, leading to poor five-year survival rates of about 25% across all stages, and as low as 8% in cases with distant recurrence [5,6]. The growing adoption of low-dose computed tomography (LDCT) screening for lung cancer has led to an increase in diagnosis of early-stage NSCLC and resulted in a stage shift, where the proportion of NSCLC patients diagnosed at stages I and II is growing, while that of later stages of disease is decreasing [7,8]. For early-stage NSCLC, surgical tumor resection remains the first line of treatment, with some stage II cases followed by adjuvant chemotherapy and/or radiation if the patient is able to tolerate it [9]. Treatment is followed up by X-ray CT imaging 3-6 months after surgery as part of surveillance imaging to monitor patients for any signs of recurrence [10]. However, despite these interventions, the rate of recurrence for early-stage NSCLC remains high (30-55%), significantly increasing the risk of mortality for these patients [11,12]. Identifying patients at high risk of recurrence following surgical resection is critical for optimizing treatment strategies and improving patient outcomes. Moreover, early detection of recurrence would enable timely intervention, and may guide the selection of patients for the most suited adjuvant therapies, thus minimizing overtreatment and unnecessary treatment related toxicities. Therefore, there is a critical need to develop prognostic markers capable of predicting risk of recurrence following surgical intervention in early-stage NSCLC.

Despite advances in early diagnosis and treatment of NSCLC, a notable absence of established markers of recurrence in NSCLC remains. Currently the prognosis for NSCLC involves the tumor-node-metastasis (TNM) staging system, which incorporates tumor size, nodal involvement, and distal metastasis to stratify patients to four stages of disease [13,14]. While the TNM system provides valuable prognostic information, it presents certain limitations. Specifically, the survival time for patients within identical tumor stages varies widely, as does their incidence of recurrence [2,15,16]. Other studies have explored the potential of imaging-based markers of recurrence, leveraging morphological and pathological information obtained from radiological imaging modalities and histologic slides. Radiomics, a field that involves the extraction of many quantitative features from medical images (such as PET, CT, and MRI), has been utilized to provide additional information for treatment strategies and to develop models for NSCLC prognosis and treatment outcomes. For example, Wang et al. constructed a predictive model that incorporated both radiomic features and clinical variables to assess the risk of postoperative recurrence in NSCLC patients. Their results demonstrated that combining CT radiomics with clinical features achieved superior predictive performance compared to models based solely on radiomic of clinical variables. [17]. Predictive modeling has also been developed by harnessing morphological features extracted from hematoxylin and eosin (H&E) stained slides. Recent studies have explored the use of digital pathology and image analysis techniques to extract quantitative features from tumor histological slides. One notable study investigated the predictive value of models involving computerized extraction of nuclear and tumoral features from H&E slides, and found that the combination of nuclear shape, texture and architectural features successfully predicted recurrence in early-stage NSCLC patients [18]. While these studies and several others [19–21] have demonstrated the potential of imaging-based predictive models for NSCLC, there are important limitations to address. Traditional medical imaging modalities, such as CT, provide valuable structural information but overall possess low spatiotemporal resolution and lack the ability to capture subcellular functional changes that occur within tumors. The heterogeneity of NSCLC at the subcellular level, including variations in cellular metabolism, may have significant implications for tumor behavior and risk of recurrence [22]. These functional and molecular changes typically precede morphological changes and necessitate imaging modalities capable of detecting such dynamics [23].

Two-photon microscopy emerges as a promising tool in this regard, offering non-invasive and high-resolution imaging capabilities to probe cellular function within tissue, and potentially provide markers of recurrence in NSCLC [24].Two-photon microscopy has an intrinsic depth sectioning ability and can penetrate deeper into tissue using near-infrared excitation wavelengths [25,26]. Two-photon excited fluorescence (TPEF) from nicotinamide and flavin adenine dinucleotides (NADH and FAD, respectively) can been quantified to determine an optical redox ratio (ORR) of FAD/(NADH+FAD), which has been shown to be highly correlated with the ratio of intracellular cofactor concentrations NAD^+^/(NADH+NAD^+^) [27]. The ORR has been used in vitro and in vivo to track metabolic changes during cancer progression and characterize early stages of cancer development [28–31]. These imaging studies have demonstrated that proliferative cells are associated with a lower redox ratio indicative of an increased rate of glucose catabolism. The ORR has also been used to quantitatively assess response to therapy in organoids derived from various cancer types, including breast, colon, and pancreas, identifying heterogenous cellular subpopulations driving varied treatment-response [32–34]. Other studies [35,36], including work from our lab [37], have demonstrated that highly invasive and metastatic cancer cells resort to mitochondrial oxidative metabolism as indicated by an increase in the redox ratio. In addition to TPEF, fluorescence lifetime imaging microscopy (FLIM) is another technique that is sensitive to metabolic changes in the tumor microenvironment. FLIM measures the time a fluorophore remains in the excited state and is sensitive to changes in the molecular microenvironment, such as pH, temperature, viscosity, and protein-binding status [38,39]. FLIM has been used to investigate cellular metabolism by discriminating between the free and protein-bound states of NADH and establishing the relative amounts of these species to characterize biological samples [38,40]. For example, decreases in the protein-bound NADH lifetime were reported in pre-cancerous epithelial cells compared to normal epithelium, consistent with increased rates of glycolysis [41]. Similar findings were reported in a study involving human lung cancer samples, where the average fluorescence lifetime for NADH in the cancerous lung tissue was significantly shorter compared to normal tissue [42].

While *in vivo* optical measurements are preferred, and can be more easily performed on accessible tissues, practical constraints limit their feasibility, especially for measurements of more inaccessible tissues such as the lung. TPEF optical endpoints have previously been reported in *ex vivo* tissues, elucidating differences between normal and neoplastic lung [42,43], diabetic and non-diabetic wounds [44], and HIV-1 infected tissue sections [45]. Jones et al. demonstrated that although the ORR from *ex vivo* skin wound frozen sections was significantly higher than that of *in vivo* measurements, the metabolic trend observed between groups was preserved [46]. Walsh et al reported similar findings when comparing optical endpoints between fresh and frozen-thawed hamster cheek pouch tissues, attributing the increase in ORR of frozen-thawed tissue to both a decrease in NADH fluorescence intensity and increase in FAD fluorescence intensity [47]. While not ideal, *ex vivo* frozen tissue sections provide unique access to otherwise inaccessible tumors, and optical imaging of these tissues still provide us with unique insight into underlying mechanisms of disease.

The overall goal of this study was to investigate the optical changes associated with long term outcome in treatment-naïve NSCLC tumors, and whether these optical endpoints can serve as predictive markers of recurrence in NSCLC. To do this we utilized high-resolution optical microscopy and image processing techniques to generate quantitative markers of metabolic activity in cryosections of primary tumors. We found that NSCLC tumors with metastatic recurrence presented with increased ORR compared to those experiencing no future recurrence. Additionally, to test the predictive capabilities of these endpoints, three predictive classifiers based on logistic regression, SVMs, and KNNs were developed by utilizing optical endpoints and clinicopathological patient feature datasets and revealed that the SVM model built with a combination of optical markers (AUC = 0.74) was capable of outperforming clinically relevant features in predicting risk of recurrence in NSCLC (AUC = 0.62). Moreover, by training a deep learning artificial neural network using whole-tissue section optical image data, we show that predictive performance outperforms that of clinical features and summary optical endpoints (AUC =), demonstrating the feasibility of using a CNN-based approach to predict risk of recurrence in early-stage NSCLC.

## Methods

### Study Design

Twenty-five NSCLC specimens were obtained from the Lung Cancer Biospecimen Resource Network (LCBRN) and the Cooperative Human Tissue Network (CHTN). These specimens consisted of a mixture of adenocarcinoma and squamous cell carcinoma at stage II of disease and were resected from the lung prior to the patient receiving any type of therapy (Figure 1A). For each tissue, a specimen and patient detail report was made available, listing follow-up information such as type of adjuvant therapy later received, metastatic status, and response to therapy. All patients in this cohort received adjuvant therapy, involving a combination of radiation and cisplatin and/or other chemotherapy drugs. The clinical detail reports were used to evaluate the follow-up data of each patient and classify them into groups that reflected their eventual outcomes (Figure 1B). Patients were classified into the metastatic group (N=11) if they presented with metastatic lesions throughout their follow-up and classified into the non-metastatic group (N=14) if they presented with no metastasis (Table 1). Similarly, patients were classified as responders if presenting with no evidence of disease after their second year of follow-up. Conversely, if the patient presented with tumor recurrence, they were classified as non-responders. All specimens had been previously flash frozen in liquid nitrogen and upon receipt were sectioned in a Leica CM1860 cryostat (Wetzlar, Germany) into sections 30 µm thick and placed in −80°C for storage until time of imaging (Figure 1C).

**Figure 1.**
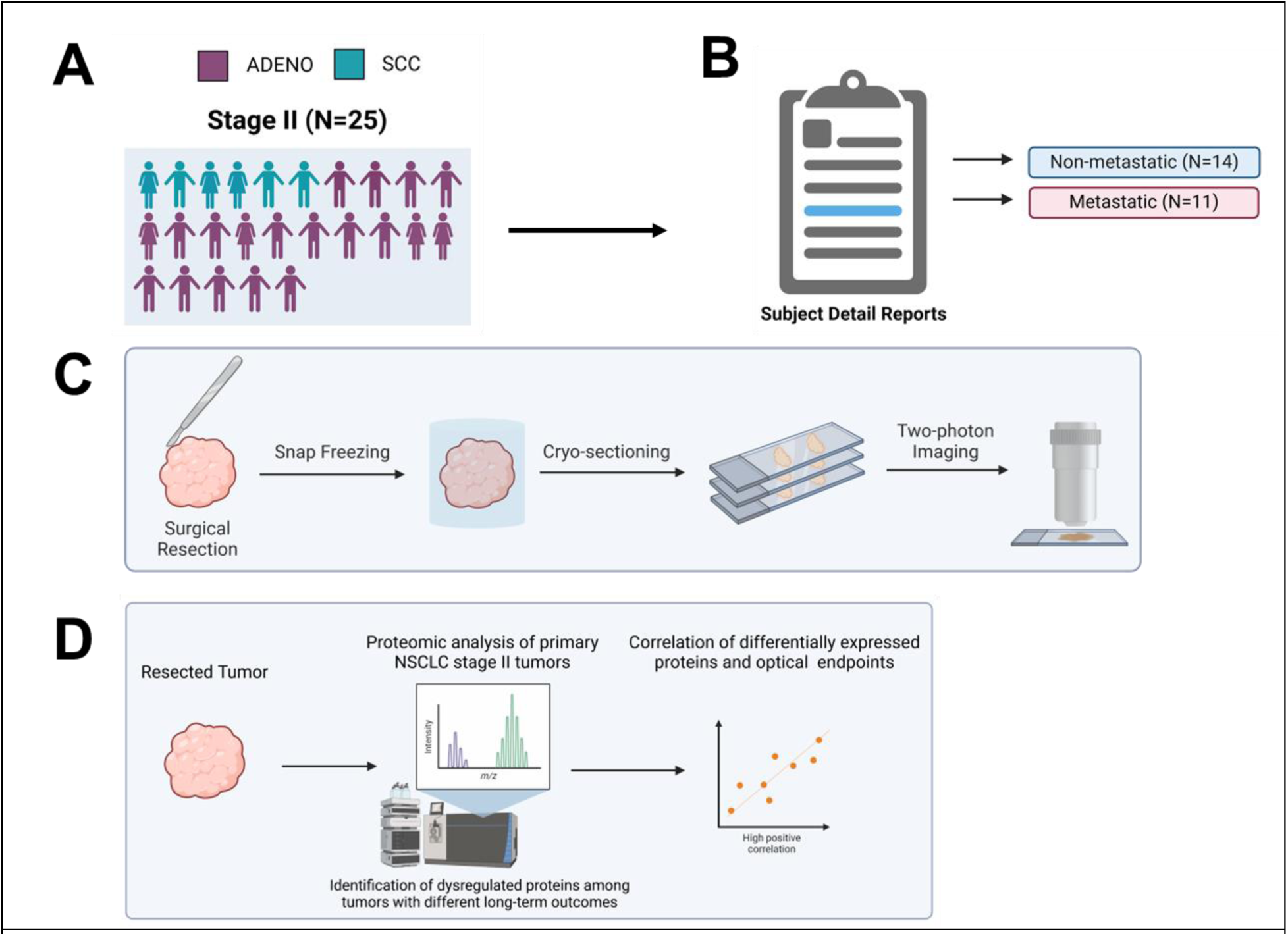
Overview of study design. Twenty-five (25) NSCLC samples were collected prior to treatment via primary tumor resection, and separated into metastatic and non-metastatic groups based on subject-detail reports (**A,B**). Upon receipt of flash-frozen samples, they were sectioned and mounted on slides for optical imaging (**C**). Tumor not used for imaging was subjected to proteomics analyses to identify dysregulated protein expression between groups (**D**).

**Table 1.** Clinical characteristics of NSCLC patients.

|  | Clinical Outcome (patient numbers) |  |  |  |
| --- | --- | --- | --- | --- |
|  | Non-Responder | Responder | Metastatic | Non-metastatic |
| Diagnosis |  |  |  |  |
| Adenocarcinoma | 12 | 7 | 10 | 9 |
| SCC | 2 | 4 | 1 | 5 |
| Stage |  |  |  |  |
| II | 14 | 11 | 11 | 14 |

#### Histopathology

Upon completion of two-photon imaging, NSCLC tissue sections were subjected to hematoxylin and eosin (H&E) staining and Verhoeff Van Gieson (VVG) elastin staining. H&E staining provided general morphological information, allowing the identification of various tissue components, namely tumor cells, necrotic and stromal regions. These regions were identified and characterized with the help of an expert pathologist at the University of Arkansas for Medical Sciences. VVG staining specifically highlighted elastin fibers as well as collagen within the tissue. A Hamamatsu NanoZoomer-SQ digital slide scanner was used to provide whole-slide histology images that were then used as reference images for the segmentation process of TPEF images (Supplementary Figure 1).

#### Acquisition of 755/855 nm two-photon excited fluorescence (TPEF)

Two-photon microscopy was used according to established methods previously described [89]. Imaging was performed using a two-photon microscope consisting of a Bruker Ultima Investigator Series laser scanning microscope (Middleton, Wisconsin), and an Ultrafast Ti:Sapphire laser (Insight X3, HP, Spectra Physics, Inc.). Samples were imaged (512×512 pixels; 13-bit depth; 0.876 pixels/µm; 1.6 µs pixel dwell time) with a 20x, 1.0 NA water-immersion objective (Olympus; Tokyo, Japan). The laser excitation source was tuned to 755 nm and 855 nm to collect NADH and FAD TPEF, respectively. Images were acquired using GaAsP photomultiplier tubes (PMTs) with a 460/20 nm bandpass filter for the 755 nm excitation, and a 525/50 nm filter for 855nm excitation. Three to five regions of interest were imaged per sample.

#### Fluorescence lifetime imaging (FLIM) at 755 nm

NADH FLIM images were acquired from the same regions of interest as TPEF. FLIM images were processed using SPCImage 8.5 (Becker & Hickl Gmbh; Berlin, Germany). Images were collected over an integration time of 2 min (pixel dwell time of 0.8 µs) and lifetime decay curves were fit to a bi-exponential model to ascertain the free and long lifetime components.

#### Image processing and quantitative analysis

Autofluorescence intensity images were normalized to laser power and PMT gain calibrated to µM concentrations of fluorescein (Figure 2A) [27]. Manual-traced masks were generated by using histological (H&E and VVG stains) images as references to isolate tumor cells from elastin-rich and necrotic regions within TPEF images from NSCLC tissue sections. In addition to manually-traced masks, Otsu’s method was applied in a custom MATLAB script to create inverse masks that specifically highlighted areas of elastin and necrotic regions within TPEF images (Figure 2B). These masks were combined to effectively isolate tumor cells of interest and the resulting mask was applied to the corresponding redox ratio and FLIM images. A pixel-wise optical redox ratio [FAD/(NADH+FAD)] was calculated with previously normalized TPEF images, and an average of the redox ratio for each field of view (as well as an average for all FLIM parameters) was calculated from ORRs located only within the masked regions (Figure 2C, D).

**Figure 2.**
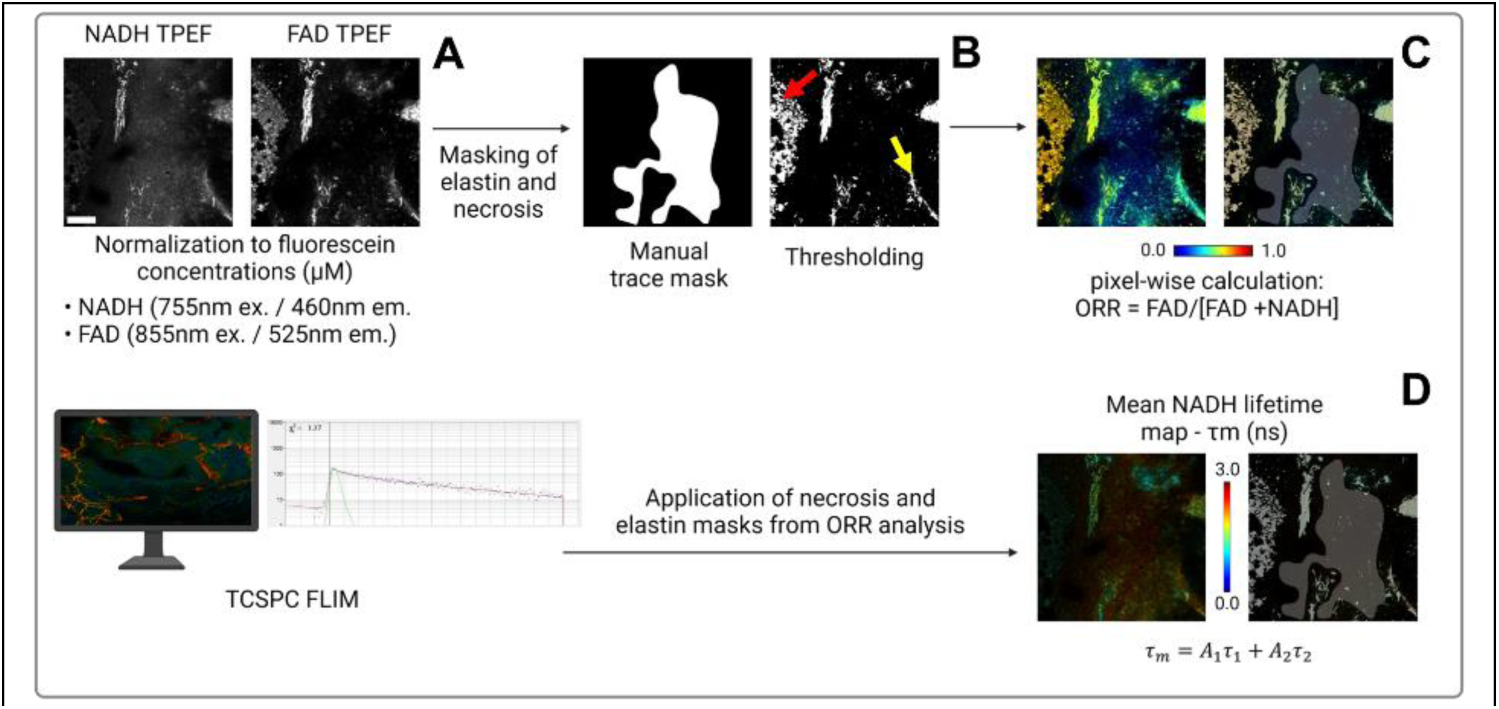
Image processing workflow for TPEF imaging and FLIM. Raw two-photon excited fluorescence intensity images (**A**) were processed with manual trace masks to exclude elastin (yellow arrow) and necrosis (red arrow) contributions (**B**). Pixel-wise calculations of the optical redox ratio were generated (**C**). Masks previously generated for ORR maps were also applied to NADH FLIM images (**D**).

#### Proteomics of NSCLC samples

Proteomic analyses and bioinformatics of the human NSCLC samples was performed at the IDeA National Resource for Quantitative Proteomics (Little Rock, AR) to determine changes in metabolic pathways to correlate to and validate the optical metabolic endpoints (Figure 1D). In brief, 150 µg of whole cell lysate protein from each tumor was trypsin digested using the FASP procedure. UPLC off-line peptide fractionation with basic pH reverse-phase chromatography (bRPLC) was used to fractionate peptides into 40 fractions. Each fraction was resolved by in-line acidic pH reverse-phase chromatography with a nanoACQUITY UPLC system coupled to a Thermo Orbitrap Eclipse mass spectrometer. High resolution MS and MS/MS data was collected. Proteins were identified with MaxQuant (<1% FDR). A label-free intensity-based quantitation approach termed iBAQ was used [48,49] to determine the differentiating levels of proteins between groups. Log2-transformed iBAQ data was analyzed using 2-sample t-tests to conduct pairwise comparisons among the groups. Pairwise differences in group means were calculated along with unadjusted P values. Unadjusted P values were converted to q-values using the FDR procedure [50]. A protein was declared to be differentially expressed if (a) any 2 of its 3-group means were different by ≥1 log2 units (a ≥2x change) and (b) the difference in question had an FDR q≤0.05. Pathway analyses were carried out using IPA. Metabolically relevant pathways were explored and proteins that are differentially expressed within these pathways were identified within each treatment group.

### Statistical analysis

To identify statistically significant differences in the ORR and FLIM parameters for both metastatic and non-metastatic groups, a mixed model analysis was performed with metastatic status set as a fixed effect and subject nested within group set as a random effect. All stats analyses for these comparisons were run using JMP® Pro 17. Correlation between ORR and KEAP1 expression was determined using Pearson’s correlation in GraphPad, as well as Kaplan-Meier survival curves.

### Supervised machine learning approaches and data preparation

We evaluated the predictive performance of three different machine learning models (logistic regression, SVM, and KNN) across three feature sets: (1) optical endpoints consisting of the mean ORR and NADH intensity, (2) clinicopathological features consisting of age and cancer subtype (ADENO/SCC), and (3) a combination of the optical and clinical features.

We conducted a retrospective analysis on the NSCLC 25 patient cohort to evaluate the predictive performance of three different machine learning predictive models - Logistic Regression, Support Verctor Machine (SVM), and k-Nearest Neighbors (KNN) - in discriminating between tumors that did and did not recur. To do this, we created three different feature sets: (1) optical endpoints consisting of the mean ORR and NADH intensity, (2) clinicopathological features consisting of age and cancer histological subtype (ADENO/SCC), and (3) a combination of the optical and clinical features. Categorical variables within the clinical feature set included cancer histological subtype and was encoded as a binary value. Features were normalized to improve model convergence and comparability, and leave-one-out cross validation (LOOCV) to assess model performance. The Area Under the Receiver Operating Characteristic Curve (AUC) was computed and compared among models.

## Results

### Optical redox ratio is significantly higher in treatment-naïve NSCLC that had metastatic recurrence

Figures 3A and 3B present representative NADH and FAD intensity images from the non-metastatic and metastatic groups. Both tumors shown here are adenocarcinomas. Elastin structures and necrotic areas were avoided as best as possible during imaging, and during image processing, any additional elastin/necrotic areas present in the images were masked out and not included in the calculation of the optical redox ratio (Fig. 3C, 3D). We observed a statistically significant increase in the average optical redox ratio (p=0.017) of the metastatic group compared with the non-metastatic group (Fig. 4E). This finding is consistent with previous work from our lab revealing increases in the redox ratio with increasing metastatic potential in cells, as well as other studies from other labs highlighting a decrease in NADH concentration in metastatic cancer [35,51].

**Figure 3.**
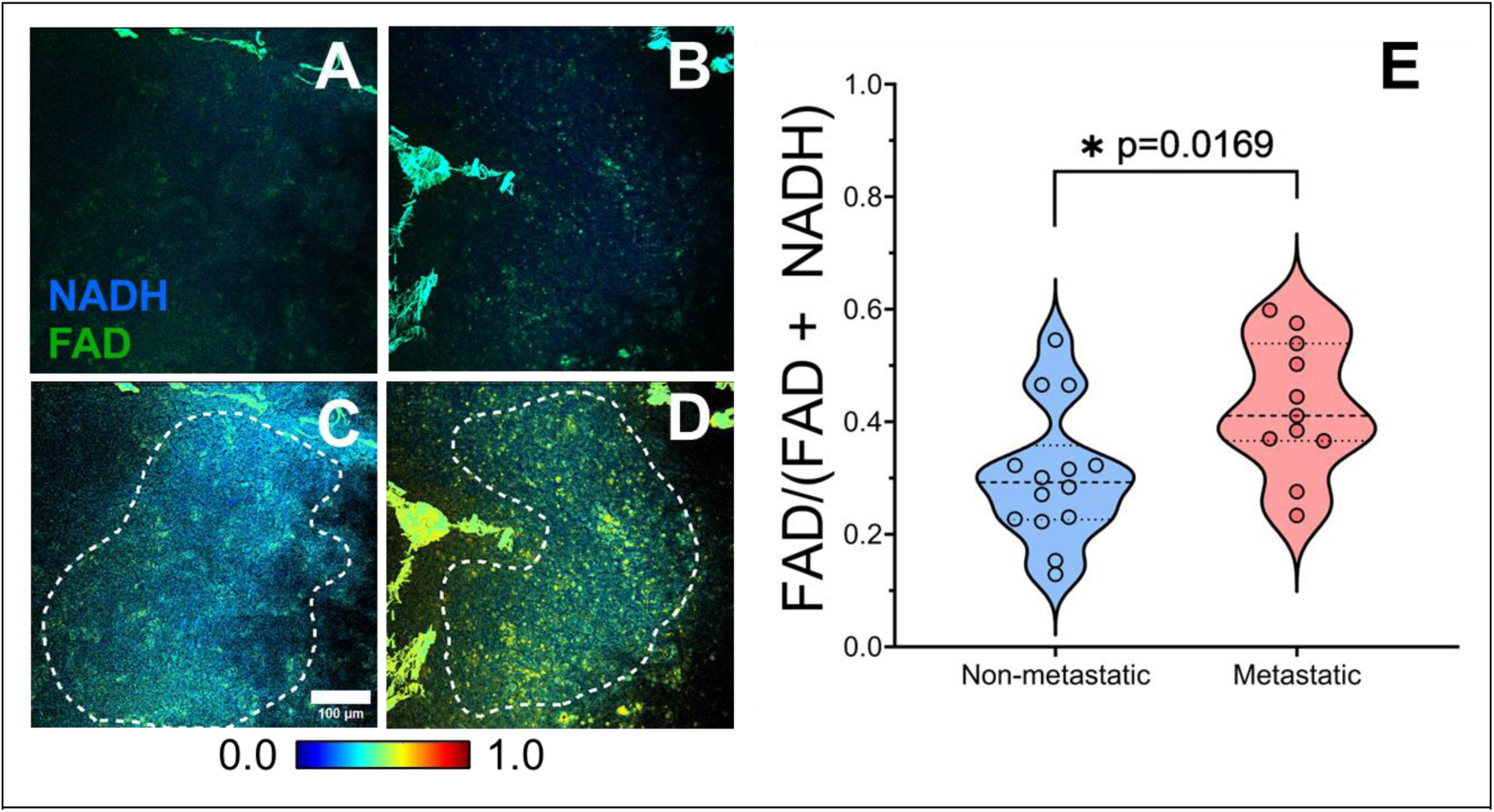
Normalized intensity images displaying NADH (blue) and FAD (green) signal (**A**, **B**) for representative ORR maps from non-metastatic (**C**) vs. metastatic tumors (**D**). A statistically significant increase in the optical redox ratio was observed in the metastatic tumor group when compared to non-metastatic (**E**). Each data point represents a patient. Scale bar: 100 µm.

**Figure 4.**
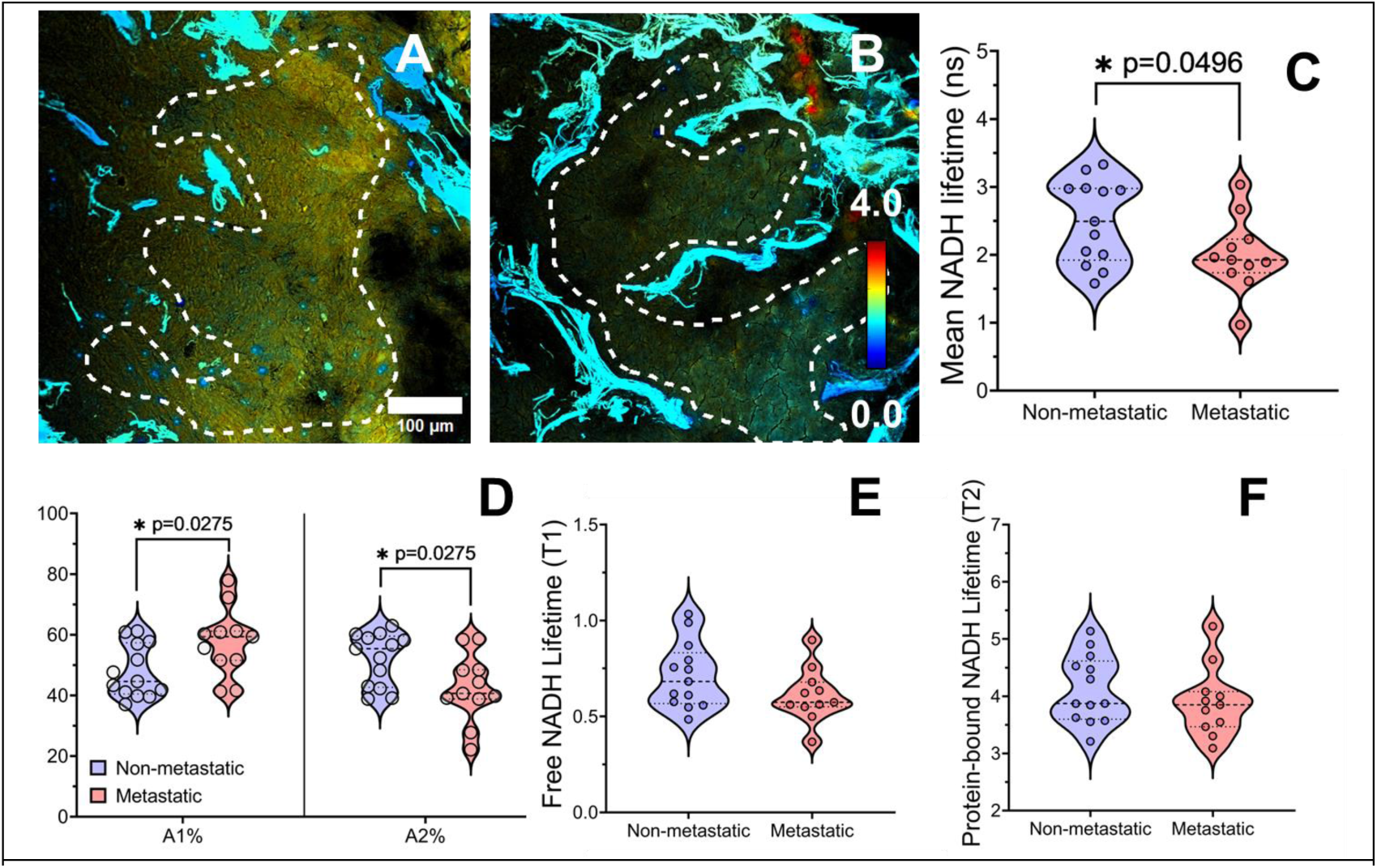
Representative mean NADH lifetime maps from non-metastatic (**A**) vs. metastatic tumors (**B**). Quantification of the average mean NADH lifetime and other lifetime parameters revealed no significant differences between metastatic and non-metastatic tumors (C-F). Each data point represents a patient. Scale bar: 100 µm.

To further characterize changes in the metabolic profiles of the NSCLC samples, we also analyzed the FLIM images (Fig. 4A, B). NADH presents very different lifetimes depending on whether it is free or protein-bound [40]. Bi-exponential decay fit models were used to separate the short (A1) and long (A2) lifetime components of free and bound NADH, respectively. A1 and A2 are the respective relative contributions to the total fluorescence. We found a significantly shorter mean NADH lifetime (τ_m_) and fraction of protein-bound NADH in tumors that eventually experienced metastasis (Figure 4C, D). Together, these results demonstrate that there are measurable differences in multiple metabolic imaging-based endpoints of NSCLC tumors based on their eventual outcome that can be exploited to evaluate long-term outcome in treatment-naïve tumors.

### High ORRs in primary NSCLC tumors are correlated with low levels of KEAP1

Following proteomic analyses, key proteins such as the kelch-like ECH-associated protein 1 (KEAP1) were identified as differentially expressed in the responder vs non-responder tumors (Supplementary Figure 2). The KEAP1-nuclear factor erythroid-2-related factor-2 (Nrf2) pathway plays a key role in protecting cells from oxidative stress and has been linked to resistance to chemotherapy, radiation, and targeted therapies in NSCLC [52]. Considering the dysregulation in KEAP1 expression in our cohort of NSCLC tumors, and the role KEAP1 plays in metabolic rewiring, we decided to investigate the association of KEAP1 expression with the ORR. A significant negative correlation between the mean redox ratio for all NSCLC samples and their corresponding KEAP1 proteomic scores was found (Figure 5A; p = 0.0371, R = −0.4276). Previous studies have shown how low or absent KEAP1 expression is linked to a poorer overall survival in NSCLC [53]. Additionally, Kaplan-Meier survival curves for our 25 NSCLC patient cohort revealed that patients with high ORRs had a significantly greater risk of tumor recurrence (Figure 5B). Altogether, these findings further highlight the clinical potential of optical metabolic imaging to establish markers of recurrence in early-stage NSCLC.

**Figure 5.**
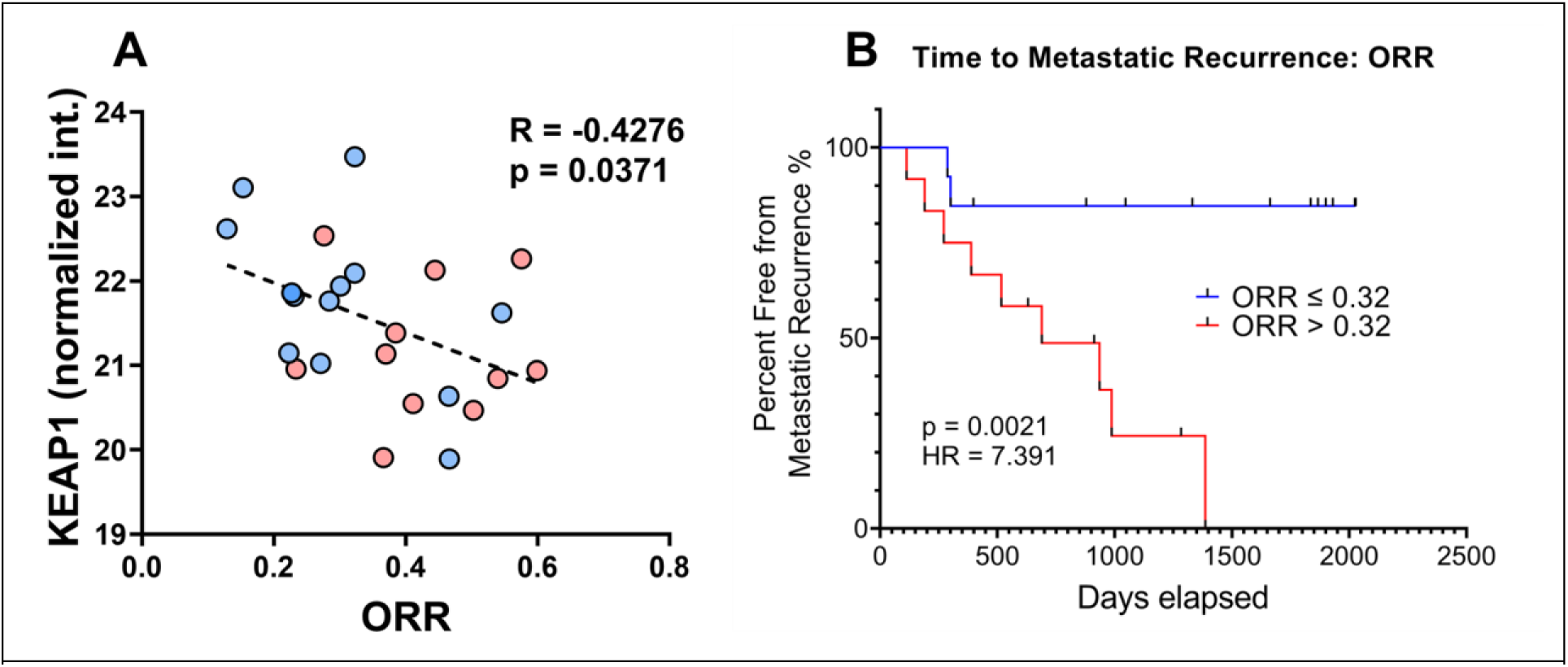
High ORR is significantly correlated (p = 0.03) with low Keap1 expression across NSCLC patients (**A**). Kaplan-Meier survival curves demonstrate greater risk of recurrence in patients with high optical redox ratios (ORR>0.32) (**B**).

### Predictive modeling based on optical metabolic features outperforms clinicopathological features for identifying tumors that did and did not metastasize

Following the evaluation of the predictive performance of three different machine learning models (logistic regression, SVM, and KNN) across three feature sets, we found that overall SVM modeling outperformed logistic regression and KNN. Moreover, when considering the different feature sets, SVM modeling resulted in a higher AUC when trained on optical features alone (AUC = 0.74) compared to clinical parameters alone (AUC = 0.62) (Figure 6). These findings illustrate the potential of optical biomarkers to outperform some of the most clinically relevant features in determining risk recurrence in NSCLC.

**Figure 6.**
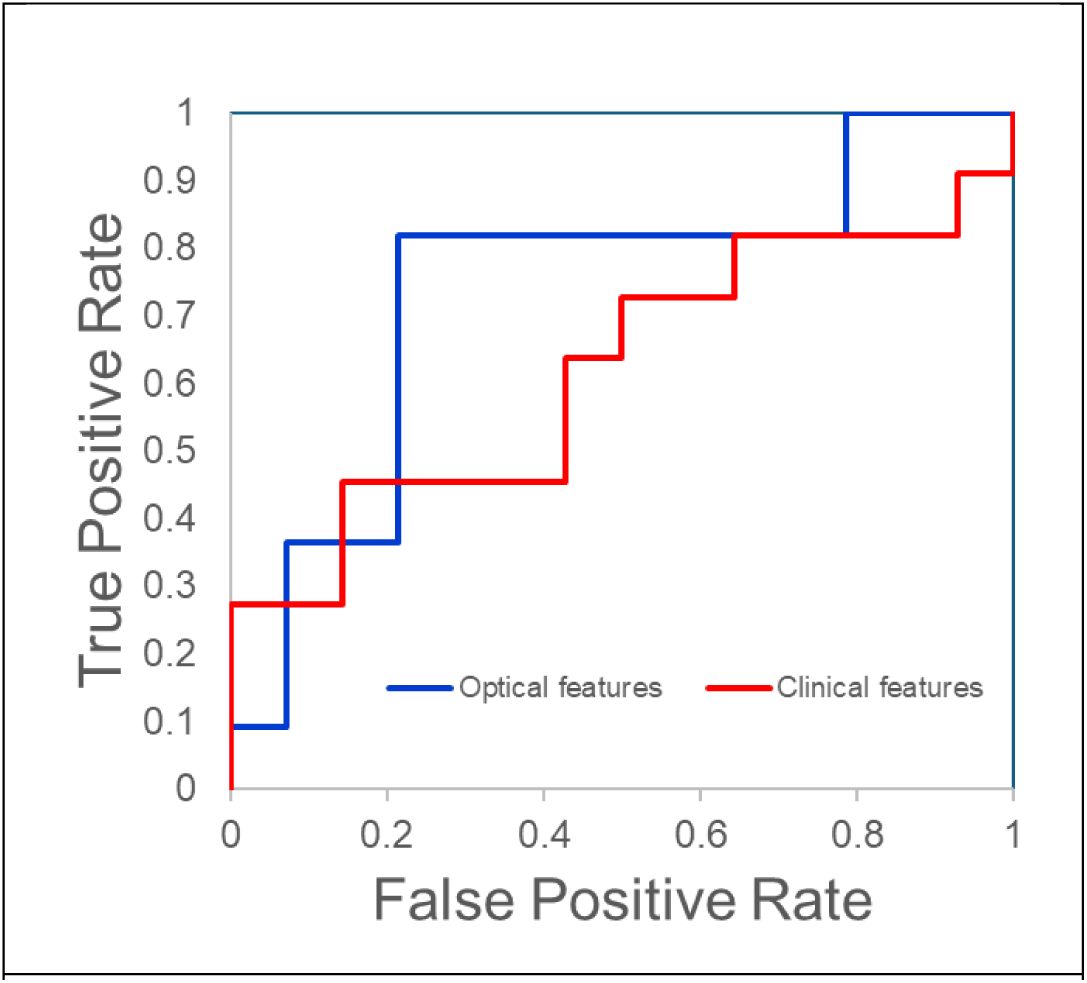
ROC curves comparing the performance of optical parameters alone (ORR and NADH intensity) to clinicopathological features alone (age and cancer subtype) using a support vector machine (SVM) model. AUC for optical parameters (0.74) is greater than clinical parameters (0.62).

## Discussion

Two-photon imaging of cellular metabolism has been used in various contexts, such as metabolic changes during cancer progression, differentiation between normal and cancerous tissue, and monitoring stem cell differentiation [27,30,31]. In addition, researchers have also used optical metabolic imaging of tumor organoids to investigate treatment response of cancers of the pancreas, head and neck, and breast [54–56]. Specifically, previous studies have linked increases in the ORR with poor prognosis. Using a human melanoma mouse xenograft model with high and low metastatic risk, Peng et al. found that the more invasive and metastatic tumors were more oxidized compared to the indolent and less metastatic ones, presenting with an increase in ORR and lower NADH [57]. Within the context of human NSCLC, optical imaging has mainly been utilized for characterization and quantification of the structural makeup of NSCLC, and for diagnostic purposes, in differentiating lung tumor from surrounding healthy tissue [42,43,58,59]. Wang et al. utilized a combination of TPEF and second-harmonic generation (SHG) signals to differentiate cancerous from non-cancerous frozen-thawed lung tissues [58]. Jain et al. also used TPM to characterize histopathological features in fresh NSCLC and conducted a blinded diagnostic analysis using TPM images, presenting both TPEF and SHG signals, that successfully differentiated neoplastic from nonneoplastic lung tissue [60]. Using a compact, portable microscope, Huizen et al. combined third and second harmonic generation (THG/SHG) and TPEF signals for the rapid identification and characterization of tissue architecture and cellular morphology in *ex vivo* fresh lung tissue. They found that the THG/SHG/TPEF images presented more information when compared to standard histopathology, positioning THG/SHG/TPEF as a promising tool for clinical intraoperative evaluation of NSCLC tissue [61]. In non-human lung tissues, Pavlova et al. utilized TPM to investigate a murine model of lung cancer and reported distinct morphological and spectral differences between *ex vivo* neoplastic lung and normal [43].

In comparison to previous studies utilizing TPM to investigate normal and NSCLC tissues, we report on a tumor-to-tumor comparison, aiming to determine whether optical markers from TPM can distinguish tumors based on their eventual metastatic outcomes. We employed image masks that isolate tumor cells within all experimental groups and remove elastin and necrotic areas to eliminate the confounding effects of these components. This approach provides a more accurate and straightforward assessment of the optical characteristics of tumor cells within the NSCLC environment.

The higher ORR observed in the metastatic tumor group proves consistent with previous work from our lab revealing increases in the ORR with increasing metastatic potential in cells, as well as other studies reporting a decrease in NADH concentration in metastatic cancers [35,51]. Metastatic and non-metastatic tumors also exhibited statistically significant differences in FLIM parameters, as well as notable variations in their data distributions as depicted by the violin plots (Fig 4). These distributions point to inherent heterogeneity and complexity within these tumors. This variability may be due to a multitude of factors, including inter and intra-tumoral biological variability, and the highly heterogeneous metabolic landscape of NSCLC wherein both enhanced glycolysis and oxidative phosphorylation have been reported on within individual tumors[22].

In addition to optical imaging, proteomic analyses were conducted, revealing dysregulated expression of KEAP1 in NSCLC tumors with poor long-term outcome. In our hands, we observed that low KEAP1, which is associated with poor long-term outcome, overlapped exclusively with regions of high ORR, which we have found in metastatic cancer cells. Gong et al showed how activation of the KEAP1/NRF2 pathway decreases intracellular ROS levels [62]. Their results revealed significantly decreased ROS levels in KEAP1-mutated lung cancer cells, leading them to conclude that KEAP1 loss is linked to increased resistance of LC cells to oxidative stress. The KEAP1/NRF2 pathway has been shown to play a key role in regulating mitochondrial metabolism, and thus in cancer progression, by directly affecting the transcription of genes heavily involved in the respiratory chain complex [63]. Alterations in the KEAP1/NRF2 pathway has been associated with dysregulated metabolism in cancer, including rewiring of glucose metabolism and lipid metabolism [64–66]. In the context of mutated/depletion of KEAP1, previous studies report on NRF2 activation, leading to a highly reductive tumor microenvironment with an increased NADH/NAD^+^ ratio and downregulated ROS levels [67]. Within our context of optical endpoints, this would translate into a lower ORR. However, in a subset of five adenocarcinoma samples, we observe that lower KEAP1 is associated with a higher ORR, and high KEAP1 expression was associated with lower ORR. In previous work, we have shown that increased ROS levels in lung cancer cells following treatment with a HIF-1 inhibitor were associated with an increase in ORR [68]. Further investigation into the role of ROS within these samples is required to discern these trends, given that the interplay between ROS and cellular redox state is complex and still not fully understood [69]

Diagnosis and treatment of NSCLC can prove challenging and accurate prediction of long-term outcomes is important for guiding treatment decisions and improving patient outcomes. Predictive modeling can be used to identify important risk factors and predictors of long-term outcome in NSCLC patients. The goal of this study was to identify potential variables that aid in predicting the metastatic status of NSCLC patients by constructing supervised machine learning models that utilize optical marker and clinicopathologic features. Specifically, our results reveal how SVM modeling built with optical metabolic endpoints outperforms clinicopathological features. We recognize that although our dataset is limited in size, it presents promise for future studies involving larger patient cohorts as well as the involvement of deep learning models that utilize entire high-resolution optical image data. Larger datasets will provide for more robust training that can both increase performance and improve generalizability. Together, our results demonstrate the potential of optical markers to serve as prognostic indicators of risk of recurrence in NSCLC, even when compared against clinically relevant features such as age and histological subtype.

## Notes

### Competing Interest Statement

The authors have declared no competing interest.

### Summary of Updates

This version of the manuscript has been revised to update Results section and corresponding figures.

