## Supplemental Material for "Optical imaging of treatment-naïve human NSCLC reveals changes associated with metastatic recurrence"

**Supplementary figures:**

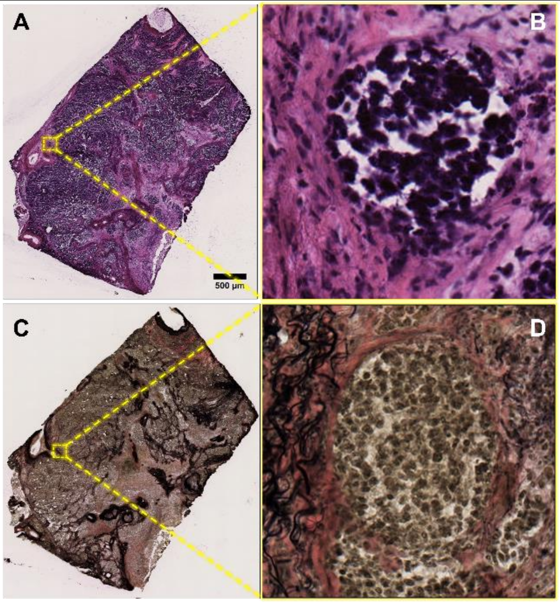


**Supplementary Figure S1.** NSCLC tumor section stained with hematoxylin and eosin (**A**) and Verhoeff-van-Gieson stain (**C**). Inset (**B**) depicts nest of tumor cells (nuclei in blue-purple in the center surrounded by stromal tissue (pink). Inset (D) depicts elastin (black) and collagen (pink) structures surrounding nest of cells. Imaged at 20X. Scale bar: 500 µm.

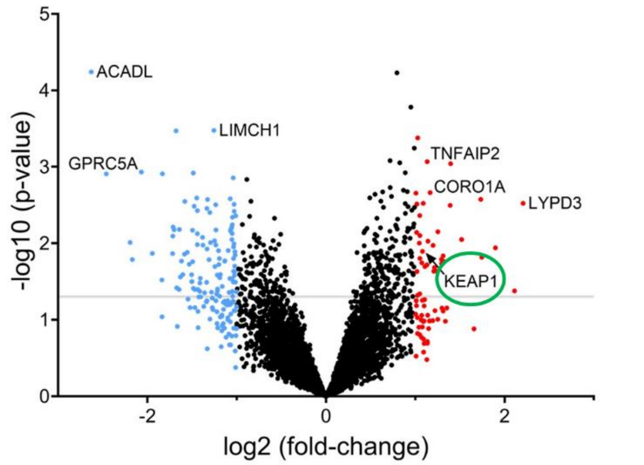


**Supplementary Figure S2.** Volcano plot displaying the log2 fold-change on the x-axis against the –log10 p-values on the y-axis. Red data points correspond to up-regulated and blue data points to down-regulated proteins. Gray horizontal line depicts cutoff for p-values (0.05). Proteins such as kelch-like ECH-associated protein 1 (KEAP1) were identified as differentially expressed in NSCLC tumors.
